# Daily rhythms in metabolic and locomotor behaviour of prematurely ageing PolgA mice

**DOI:** 10.1101/2024.03.27.586233

**Authors:** Amit Singh, Dilara Yilmaz, Esther Wehrle, Gisela A. Kuhn, Ralph Müller

## Abstract

Ageing is an inherent and intricate biological process that takes place in living organisms as time progresses. It involves the decline of multiple physiological functions, leading to body structure and overall performance modifications. The ageing process differs among individuals and is influenced by various factors, including lifestyle, environment, and genetic makeup. Metabolic changes and reduced locomotor activity are common hallmarks of ageing. Our study focuses on exploring these phenomena in prematurely ageing PolgA^(D257A/D257A)^ mice (also known as PolgA) aged 41-42 weeks, as they closely mimic human ageing. We assess parameters such as oxygen consumption (VO2), carbon dioxide production (VCO2), respiratory exchange ratio (RER), and locomotor activity using a metabolic cage for four days and comparing them with age-matched wild-type littermates (WT). Our findings revealed that VO2, VCO2, RER, locomotor activities, water intake, and feeding behaviour show a daily rhythm, aligning with roughly a 24-hour cycle. We observed that the RER was significantly increased in PolgA mice compared to WT mice during the night-time of the light-dark cycle, suggesting a shift towards a higher reliance on carbohydrate metabolism due to more food intake during the active phase. Additionally, female PolgA mice displayed a distinct phenotype with reduced walking speed, walking distance, body weight, and grip strength in comparison to male PolgA and WT mice, indicating an early sign of ageing. Taken together, our research highlights the impact of sex-specific patterns on ageing traits in PolgA mice aged 41-42 weeks, which may be attributable to human ageing phenotypes. The unique genetic composition and accelerated ageing characteristics of PolgA mice make them invaluable in ageing studies, facilitating the investigation of underlying biological mechanisms and the identification of potential therapeutic targets for age-related diseases.

## Introduction

Over the past decade, laboratory mice have emerged as a commonly used and invaluable model system to study the underlying mechanism of biological ageing in humans [1]. Several biological functions, including signalling pathways, cellular mechanisms, metabolism, and body composition, decline with ageing [2–6]. These age-related changes at the tissue and organ levels were also observed in old mice compared to young mice [7–10]. Many physiological parameters oscillate throughout the day, and the circadian clock system governs the physiological and behavioural rhythms in a 24-hour cycle. These include rest-activity cycles, metabolic rate, body temperature, food, and water intake [11–14]. At the cellular level, circadian clocks play a pivotal role in regulating daily rhythms in mitochondrial oxidative metabolism [15,16]. The mitochondria use nutrients such as lipids and carbohydrates and convert them to energy through oxidative phosphorylation, where oxygen is consumed in mitochondria, and carbon dioxide is the product of the citric acid cycle that displays daily oscillations [17–19]. The energy source used by the body in which oxygen is consumed (VO2) and carbon dioxide is produced (VCO2) is measured by calorimetry [20,21]. The ratio of VCO2/VO2 calculates the RER, which estimates the respective contributions of fats and carbohydrates to energy production, enabling the calculation of total energy expenditure [22]. The circadian rhythms, which directly influence tissue homeostasis, sleep regulation, behaviour, and metabolism, are closely linked to the ageing process [23–27]. The ageing process is marked by a reduction in mitochondrial function, which is intimately tied to energy expenditure and contributes to cellular metabolic alterations [28–31]. Previous studies have indicated that mitochondrial mutations and dysfunction are associated with ageing [32–36]. It is shown that the PolgA (D257A/D257A) mice display a premature ageing phenotype due to extensive mitochondrial dysfunction and point mutations in mitochondrial DNA [37–39]. In earlier studies, PolgA mice displayed diverse age-related phenotypes, including kyphosis and alopecia at ∼ 40-45 weeks [40]. At 40 and 46 weeks, these mice showed progressive musculoskeletal deterioration characterized by reduced bone and muscle mass and impaired function [37,38]. Additionally, dopaminergic dysfunction and behavioural and metabolic characterization were assessed in PolgA mice at 3 and 14 months, showing mitochondrial electron transport chain functions and experiencing a notable decline in activity as individuals grow older [41]. Moreover, PolgA mice have exhibited shifts in the metabolism of amino acids and lipids during 3 to 11 months of age. These alterations were examined using dynamic labelling techniques with 13C and 15N [42]. PolgA mice show distinctive indicators of musculoskeletal ageing. These mice demonstrate premature frailty and a notable bone density and muscle mass reduction, as observed in human musculoskeletal ageing processes. This resemblance makes PolgA mice a valuable model for studying and understanding the intricate mechanisms underlying age-related changes in the musculoskeletal system [43]. Additionally, PolgA mice show signs of frailty around 40 weeks, which is roughly equivalent to 20-23 months in C57BL/6 mice and 65-70 years in humans [44–46]. However, locomotor activity and metabolic behaviour indicators of premature ageing have not been thoroughly explored in these mice. Considering the crucial role of these parameters in selecting inbred strains for age-related research, this study aimed to comprehensively understand and provide normative data on locomotor activity and metabolic behaviour during the light-dark cycle. This study assesses metabolic patterns in 41–42-week-old PolgA and WT mice, focusing on variables like VO2, VCO2, RER, locomotor activity, and feeding behaviour. Our findings reveal daily rhythms in both groups, with PolgA (female) mice exhibiting a higher RER peak-to-trough ratio compared to WT mice. Additionally, female PolgA mice showed reduced body weight, walking speed, locomotor movement, and muscle grip strength compared to males and their WT counterparts. These changes were observed to be more pronounced in 41-42-week-old female PolgA mice compared to males, which have not been previously described. Therefore, PolgA mice can serve as a valuable model for investigating the mechanisms underlying the ageing process. This model allows researchers to study age-related processes and interventions within a shorter time frame, providing insights into potential therapeutic strategies for age-related disorders.

## Materials and Methods

### Animals

Our animal experiments, conducted with the approval of the local authority (Veterinäramt des Kantons Zürich, License Nr ZH35/2019), were carried out in an ethical and responsible manner. A colony of the mouse strain expressing an exonuclease deficient version of the mitochondrial DNA polymerase γ (PolgAD257A, B6.129S7(Cg)-Polgtm1Prol/J, JAX stock 017341, The Jackson Laboratory) at 41-42 weeks was bred and maintained at the ETH Phenomics Center (12h:12h light-dark cycle, maintenance food, and water ad libitum, 3-5 animals/cage). The “Kliba Nafag 3437” (Kliba Nafag, Kaiseraugst, Switzerland) maintenance diet was employed consistently throughout the study. Eight mice per sex PolgA and WT were selected for the experiment. Animals had a general health check before the experiment.

### Metabolic cage and Data collection

At the ETH Phenomics Center, the PhenoMaster system (TSE PhenoMaster, Bad Homburg, Germany) was used to examine the metabolic and behavioural characteristics of PolgA and WT mice. Each mouse was kept in a single house in a cage with ad libitum access to food and water for four days. The PhenoMaster system consists of two environmental chambers, each capable of accommodating up to eight cages and monitoring 16 mice in parallel. Each cage records physiological and behavioural measurements of parameters through sensor systems, creating a stress-free home cage environment. A calorimetry system measures carbon dioxide production and oxygen consumption in cages. The carbon dioxide and oxygen sensors were calibrated with calibration gas mixtures following the manufacturer’s recommendations. Additionally, the cages contain weight and feeding basket sensors to accurately measure body weight and food intake. The TSE system is equipped with a multi-dimensional infrared beam system that provides real-time measurement of locomotor activity in a 3D direction. The following crucial parameters related to metabolic, physiological, and behavioural parameters are recorded: i) Drinking and feeding amount, ii) Walking speed, all movement, and locomotor activities (Total movement in x and y direction in the metabolic cage and walking distance), iii) Metabolic performance VO2 consumption, VCO2 production, and RER at 10 min intervals for four days. The experiment was performed in WT and PolgA mice (male and female).

### Forelimb grip strength

The study assessed the forelimb grip strength of PolgA and their WT littermates at 41-42 weeks-old mice using a force tension apparatus (Grip Strength Meter, model 47200, Ugo Basile). Mice were positioned horizontally on a grid and pulled from their tails until they lost their grip. The force exerted was recorded, and the test was repeated five times for each mouse. The average force value (gram force) of the trials was used for analysis.

### Data analysis

Each experimental run was treated as an independent event, and the average values of each feature over time were calculated for each mouse. The data used for the analysis include WT male (n=8), WT female (n=7), PolgA male (n=8), and PolgA female (n=8) mice. Due to a technical recording issue, one WT female mouse was not included in the locomotor and metabolic parameter assessment. The first 48 hours of data were excluded from the analysis to mitigate potential stress from acclimatisation to the metabolic cage. The last 48 hours of measurements were consolidated into a single light-dark cycle by computing the average within a 24-hour window and used for the visualization. All measured metabolic and locomotory parameters are reported in Table S1. The data were further smoothed using Locally Estimated Scatterplot Smoothing (LOESS). We conducted comparative tests using analysis of variance (ANOVA) for daily rhythm of metabolic and locomotor parameters. For daily rhythm parameters, the model includes interactions among genotype, sex, and time of day. For grip strength and body weight, the model includes interactions between genotype and sex. Significance levels between the groups (e.g., male/female, day/night, PolgA/WT) were calculated using pairwise comparisons with Tukey’s post hoc correction for multiple comparisons. The p-values are reported in the figures comparing sexes in time of the day, and the estimated marginal means and contrast comparisons between the levels of interaction are provided in Table S2. The basic formula for calculating energy expenditure is typically expressed in VO2 and VCO2. Here, we used the Weir equation to find the energy expenditure EE=(3.941×VO2) + (1.106×VCO2) kcal/d. We used the dark period time (12h) average value of VO2 and VCO2 for this calculation. To study the daily rhythmic behaviour, we fit the following harmonic function to the RER data of the last 48 hours of measurement to estimate the values of the parameters *β*_0_, *β*_*k*_, *α*_*k*_. The sine function represents the vertical component of the waves, indicating the amplitude or height of the oscillations, while the cosine function represents the horizontal component, indicating the phase or timing of the oscillations. By analyzing the fitted sine curve, we calculated the periodic length of the daily rhythm, corresponding to the duration of one complete oscillation cycle.

[*y*(*t*) = *β*_0_ + *β*_*k*_cos(2π*kt*) + *α*_*k*_sin(2π*kt*)]

Where y(t) is the response variable at time t, *β*_0_ is the intercept term, *βk* are the coefficients for the cosine terms, *αk* are the coefficients for the sine terms, and 2πkt is the angular frequency (w) of the harmonic terms. Given the consistent and smooth nature of the RER data (VCO2/VO2), a harmonic function was applied to analyse the data. Meaningful conclusions were drawn based on the RER data, such as the periodic length of the daily rhythm and the peak in the night-time. Data input, cleaning, and statistical analysis were conducted using the R programming language (R Core Team 2023). ANOVA and Tukey’s post hoc test were performed using “aov” function and *Emmeans* packages in R [47]. MATLAB was used for data smoothing, fitting the harmonic model, and generating visualizations.

## Results

### Daily rhythms in oxygen consumption and carbon dioxide release

To examine and track the metabolic and locomotor behaviours, along with their daily changes, in both PolgA and WT mice (41-42 weeks old), we used PhenoMaster metabolic cages. The experimental setup is illustrated in Fig. 1A-E. The PolgA and WT mice, with eight males and females in each group, were individually housed in PhenoMaster metabolic cages. We recorded the levels of VO2, VCO2, RER, locomotor activity, water intake, and feeding patterns in 10-minute intervals over four days. Consistent with the behaviour of nocturnal animals, we observed a significant increase in VCO2 (Fig. 2A) and VO2 (Fig. 2B) during the night-time (12h) of the light-dark cycle. Further, the RER increased in both PolgA and WT mice at night-time (Fig. 3A). We observed an approximately 1.5 times increase in the VO2 and VCO2 levels at night-time compared to the daytime (12h). Female mice showed increased total VO2 and VCO2 in PolgA and WT mice (Fig. 3B, C). Notably, PolgA mice (female and male) demonstrated slightly higher RER than WT mice at night (Fig. 3D). Higher oxygen consumption is typically associated with elevated energy expenditure [48], and animals with smaller body sizes tend to have high metabolic rates, ultimately associated with high energy expenditure [49]. The RER is a widely used informative index to assess substrate utilisation in tissues, specifically the oxidation of fatty acids and carbohydrates. The RER values can vary between approximately 0.7, indicating a predominant reliance on fat as an energy source, and 1.0, indicating a preference for carbohydrates as a fuel source [50–52]. In our study, we observed that the average RER was significantly increased during the night-time of the light-dark cycle for both male and female mice, as depicted in (Fig. 3D). This finding suggests a shift towards a higher reliance on carbohydrate metabolism due to more food intake during the active phase. Moreover, when comparing the RER values between the PolgA and WT mice under both light and dark conditions, we observed a statistically significant distinction in which PolgA mice displayed elevated RER values compared to WT mice, suggesting distinct metabolic profiles between the two mouse groups during different day and night-time of the light-dark cycle. Notably, the PolgA mice show higher RER than the WT mice, signifying a shift in their fuel substrate utilisation towards a higher reliance on carbohydrates (approaching 1.0) and reduced reliance on fats as their predominant energy sources. We employed a harmonic regression model to analyse the daily rhythm of the RER, as illustrated in (Fig. 4A). In constructing the model, we used the average data from the last 48 hours of measurements, ensuring a robust representation of the cyclic pattern. The model analysis provided valuable insights into the temporal patterns of RER, showing that female PolgA mice displayed a delayed peak and peak-to-trough ratio compared to male PolgA and female WT mice (Fig. 4B). Additionally, the RER showed daily rhythmicity, contributing to the synchronised metabolic processes occurring within the 24-hour cycle. Additionally, we calculated energy expenditure during the night using the Weir equation and found that PolgA mice spend (female=6908.504, male=6529.177 Kcal/d) energy, whereas WT mice spend (female=6968.872, male=6403.492 Kcal is/d) energy. Our observation shows that female mice in WT and PolgA have higher energy expenditure than their male counterparts.

**Fig. 1.**
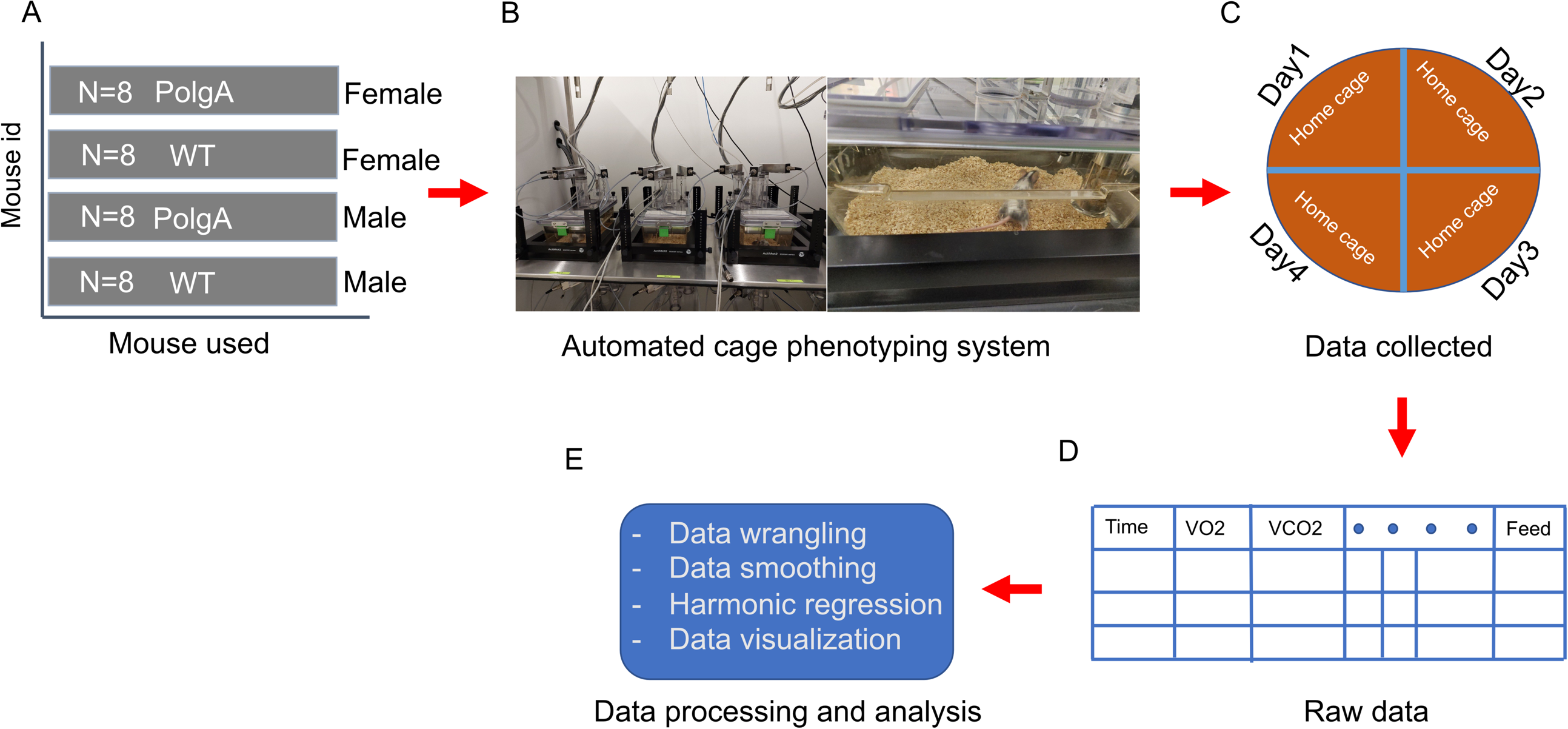
Experimental study design. (A) The study used PolgA and WT mice, with a sample size of 8 males and females per group. (B) The experimental setup included using the PhenoMaster system and an environmental chamber for automated phenotyping, which was captured in pictures. (C) The experimental measurement time for each animal was monitored for four days and nights. (D) Data were collected for each animal, where each row represented the time, and each column represented a specific measurement. (E) The collected data underwent statistical data analysis.

**Fig. 2.**
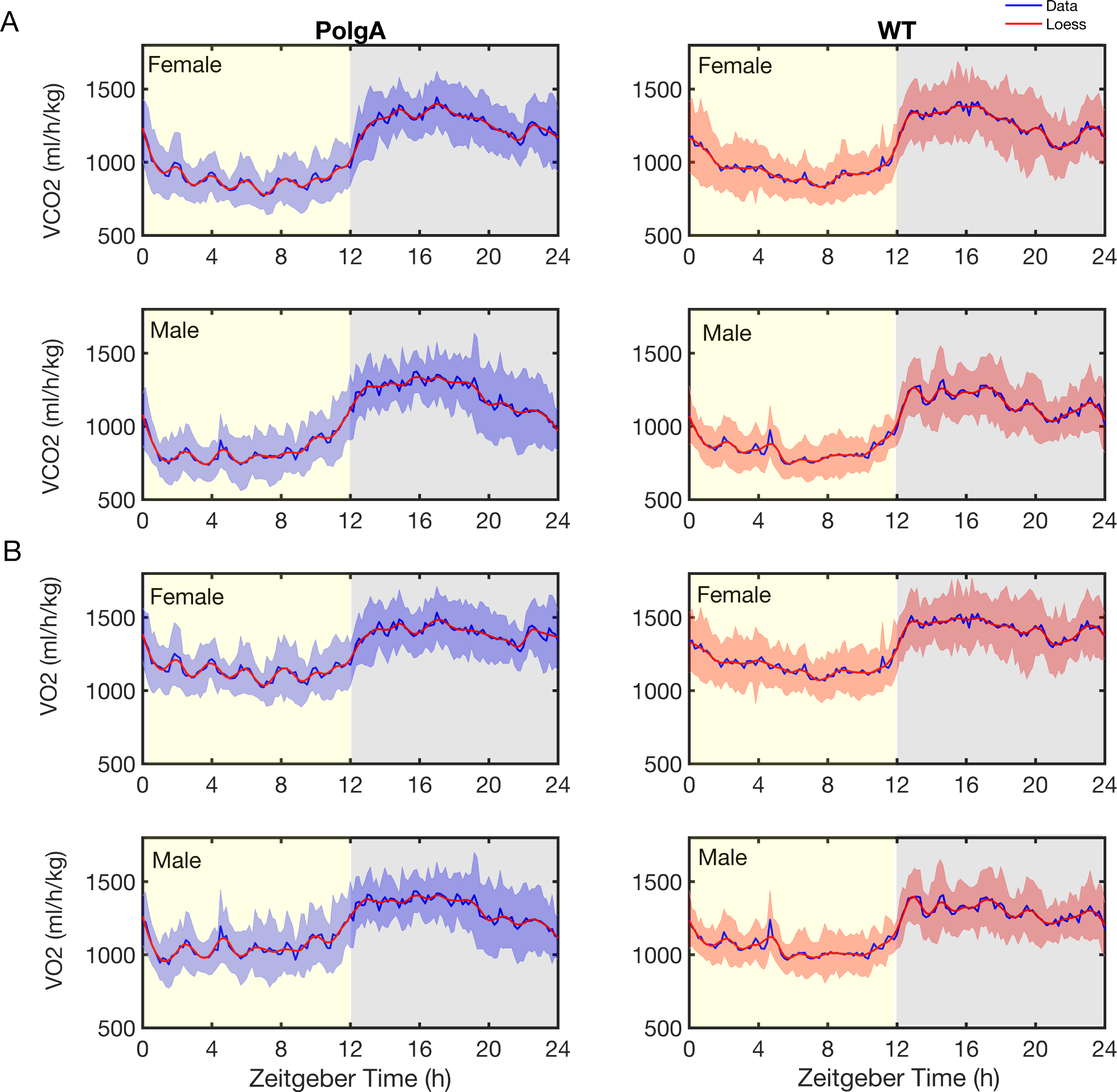
Daily rhythms in oxygen consumption and carbon dioxide release. (A) The study recorded carbon dioxide production (VCO2). (B) Oxygen consumption (VO2) was measured using metabolic cages for four days, as described in the methods section. The solid blue line represents the average values recorded at 10-minute intervals, while the blue shaded areas represent the standard deviation (SD) (WT male (n=8), WT female (n=7), PolgA male (n=8), and PolgA female (n=8)). The solid red line indicates Loess smoothing for PolgA mice in the left panel. The right panel shows the results for WT mice. The yellow shaded area indicates the 12-hour daytime, while the grey shaded area corresponds to the 12-hour nighttime.

**Fig. 3.**
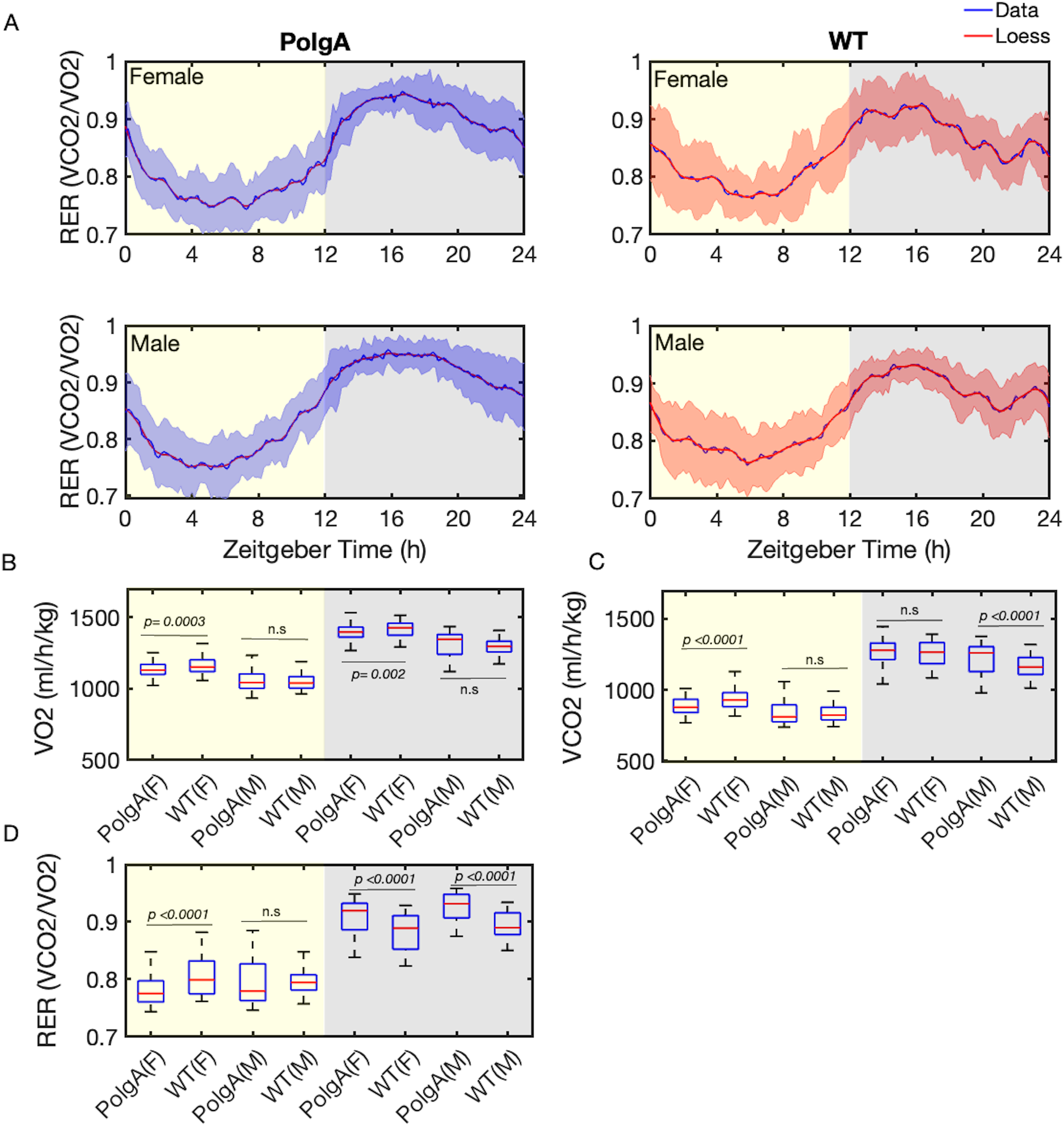
Daily rhythms in respiratory exchange ratio. (A) The respiratory exchange ratio (RER) measurements from the metabolic cage: the solid blue line represents the average value, the blue shaded areas correspond to the standard deviation (SD). The solid red line in the left panel indicates Loess smoothing for PolgA mice. The right panel displays results for WT mice, with the solid blue line representing the average value and the solid red line representing Loess smoothing, and the red-shaded areas corresponding to the standard deviation (SD). (B) The box plot of oxygen consumption (VO2) during the light-dark cycle. (C) The box plot of carbon dioxide production (VCO2) during the light-dark cycle. (D) The box plot represents the total measurement of the respiratory exchange ratio (RER). Yellow marked areas indicate the 12-hour daytime, while the grey areas correspond to the 12-hour nighttime. (B-D) The data are represented as average values of the measurements, with the central red mark indicating the median, blue whiskers representing the range, and each box representing the interquartile range (WT male (n=8), WT female (n=7), PolgA male (n=8), and PolgA female (n=8)). The figure presents the *p*-values for each comparison derived from ANOVA tests; n.s. represents no significant difference.

**Fig. 4.**
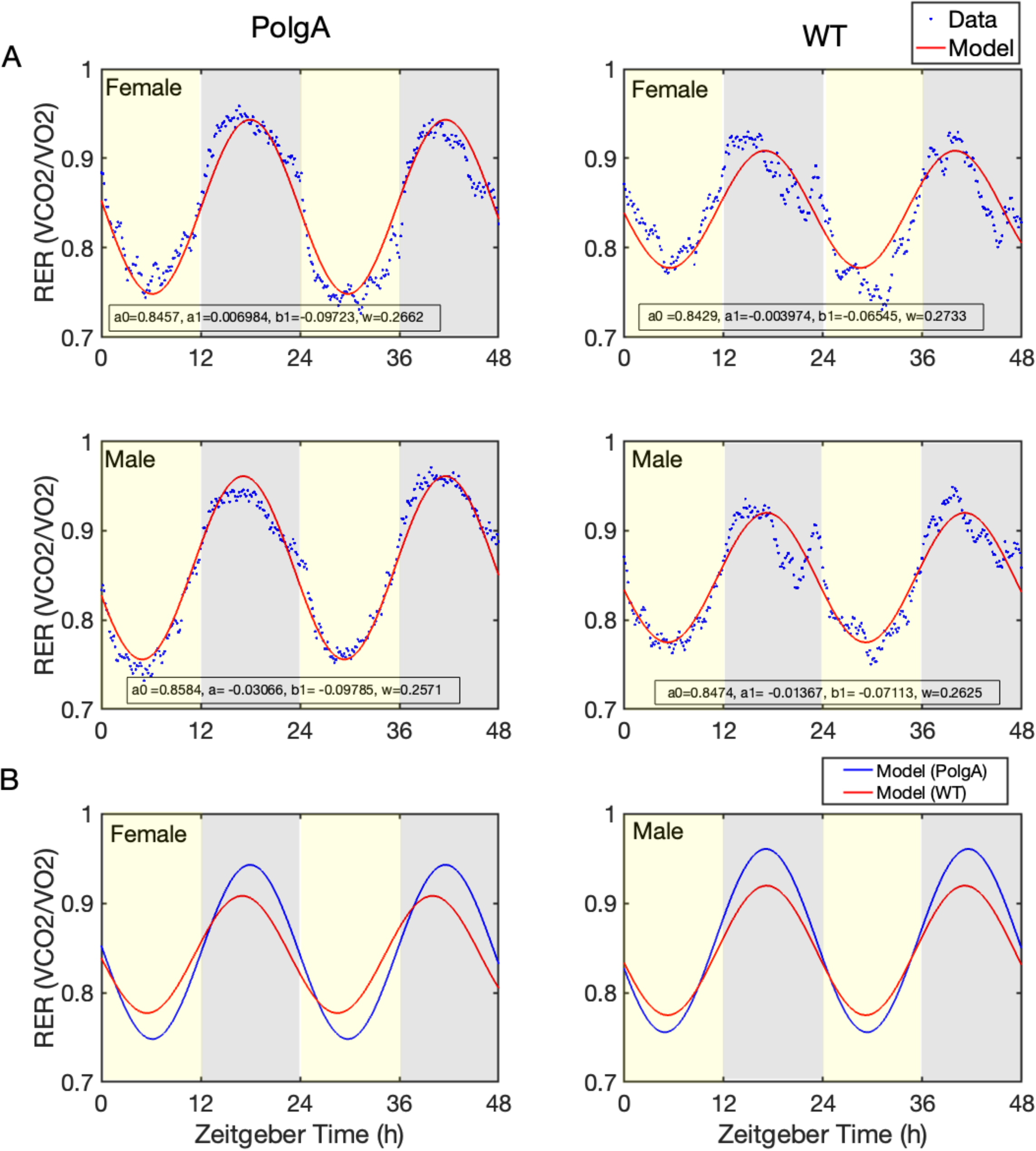
The harmonic regression models. (A) The harmonic regression model analysis was applied to respiratory exchange ratio data (RER). The solid red line represents the model prediction, and the blue dotted line represents the experimental data. The left panel of the graph shows the results for the PolgA mice, while the right panel shows the results for the WT mice. Yellow shading in the graph represents the daytime, and grey shading represents the nighttime. The model coefficients (parameters) are displayed on the plot. (B) The fitted model is represented by the blue line for PolgA and the red line for WT females (left) and males (right). The data are presented as average values (WT male (n=8), WT female (n=7), PolgA male (n=8), and PolgA female (n=8)).

### Daily fluctuations in food consumption, water intake, and body weight

We monitored these mice’s water intake and feeding habits. Assessing the patterns of food and water consumption is crucial as these behaviours serve as vital biomarkers of the mice’s overall well-being. We examined the feeding habits of these mice. Both male and female PolgA mice tended to consume more food than the WT mice at night (Fig. 5A, Fig. 5C). In contrast, male and female PolgA mice displayed similar water intake patterns (Fig. 5B). The body weight is a crucial metabolic parameter that can have significant implications for ageing and overall health [53,54]. We observed a notable reduction in the body weight of the PolgA mice in comparison to the WT mice, as illustrated in Fig. 5D.

**Fig. 5.**
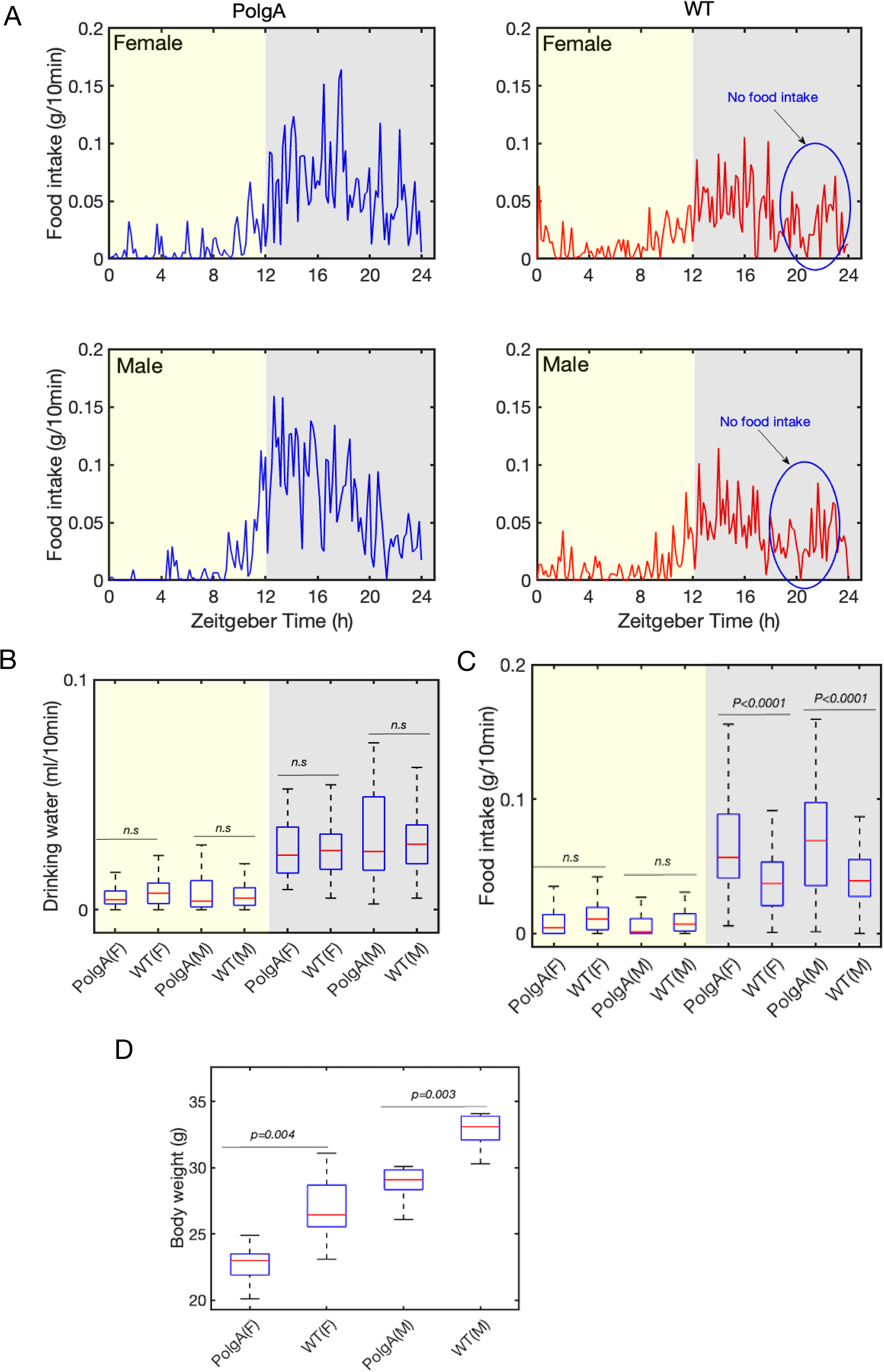
Body weight, water, and food intake. (A) The food intake during the light-dark cycle in PolgA and WT mice. The blue circle represents ‘no food intake’ during the time intervals between 20-22h. (B) The box plot represents drinking water intake during the light-dark cycle. (C) The box plot represents food intake during the light-dark cycle. The yellow area marks the daytime, and the grey-shaded area represents the dark. (B-C) The data are represented as average values of the measurements, with the central red mark indicating the median, blue whiskers representing the range, and each box representing the interquartile range (WT male (n=8), WT female (n=7), PolgA male (n=8), and PolgA female (n=8)). (D) The box plot represents the body weight of PolgA and WT mice (n=8). (B-D) The figure presents the *p*-values for each comparison derived from ANOVA tests; n.s. represents no significant difference.

### Daily locomotor activity

Investigating the locomotor activity of these mice during both day and night-time is crucial in comprehending their complete activity patterns and potential variations in their behaviour. Monitoring total movement is particularly important in the ageing process, as it fosters functional independence, maintains muscle mass and strength, and enhances cardiovascular health. In metabolic phenotyping cages, the total movement was recorded and detected through horizontal walking with beam breaks in the x and y directions. As expected, locomotor activity is elevated at night-time of the light-dark cycle. Our findings revealed that female WT and PolgA mice show significantly higher total locomotor activity than male mice (Fig. 6A, 6B) in the night-time of the light-dark cycle. Nevertheless, female PolgA mice showed reduced walking distance compared to WT mice in the night (Fig. 6C). Furthermore, the daily variations in walking speed in the PolgA mice show a substantially reduced walking speed compared to their WT counterparts at night (Fig. 6D).

**Fig. 6.**
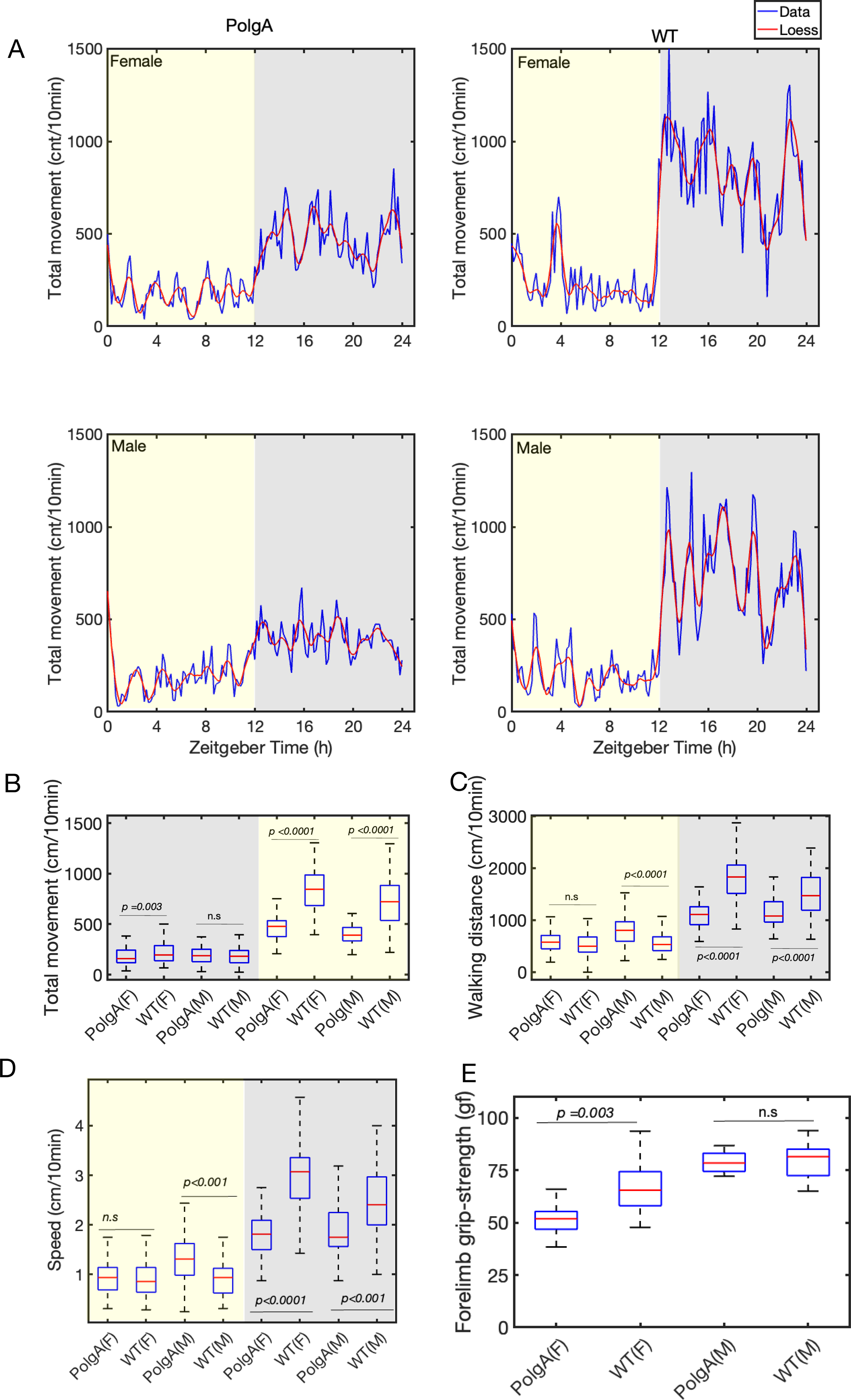
Daily rhythms in locomotor activity and grip strength measurement. (A) The daily rhythm of the total movement of both PolgA and WT mice. The solid blue line averages values recorded at 10-minute intervals. The solid red line is the Loess smoothing in the left panel for PolgA mice, and the right panel is for WT mice. (B) The box plot illustrates total movements in the x and y direction of the metabolic cage. (C) The box plot displays the total walking distance during the light-dark cycle. (D) The box plot displays the walking speed during the light-dark cycle. The yellow-marked area represents the daytime, and the grey-shaded area represents the nighttime. The left panel corresponds to females and the right one to males. (B-D) The data are represented as average values of the measurements, with the central red mark indicating the median, blue whiskers representing the range, and each box representing the interquartile range (WT male (n=8), WT female (n=7), PolgA male (n=8), and PolgA female (n=8). (E) The box plot compares grip strength between PolgA and WT mice (n=8). (B-E) The figure presents the *p*-values for each comparison derived from ANOVA tests; n.s stands for no significant difference.

### Grip strength

We assessed the grip strength of the mice, as it is a valuable measure for evaluating muscle strength. Grip strength is an essential indicator of disability, morbidity, and mortality [55]. We performed the grip strength at 41-42-week-old PolgA and WT mice. Our analysis revealed decreased grip strength in the PolgA compared to the WT mice (Fig. 6E). The observed decrease in grip strength in the female PolgA mice suggests a more pronounced premature ageing phenotype than the WT mice.

## Discussion

Ageing is associated with complex phenotypical and functional changes affecting different organs, skeletal muscle strength, energy balance, and physiological activity across the organism’s life span [54,56,57]. In recent years, ageing research has become a focal point of scientific investigation, with a particular emphasis on gaining a deeper understanding of the intricate interplay between chronological and biological ageing phenotypes. Longitudinal studies are vital for investigating ageing and pinpointing risk factors linked to age-related diseases. These studies are meticulously conducted within controlled environments to guarantee the accuracy and reliability of the findings without experimental bias across different laboratories [58–60]. In this conjuncture, we monitored oxygen consumption, carbon dioxide release, food intake, and spontaneous locomotor activity in 41–42-week-old PolgA and WT mice. Our results indicated an elevation in VO2, VCO2, and RER, levels at the night-time of the light-dark cycle (Fig. 3B-D). This is consistent with the fact that mice and rats are nocturnal animals and are most active at night-time [61]. On the other hand, circadian studies on humans or fungi show diurnal activities and are most active during the day-time [62,63]. We found a slight increase in the average RER mean value during the night-time in PolgA (female 0.90, male 0.92) due to increased food intake in comparison to WT mice (female 0.88, male 0.89) (Fig. 3D). In the context of ageing, a previous study conducted with male mice unveiled a significant decrease in their RER as they progressed in age. This decline signifies a transition towards a greater emphasis on fat metabolism [64]. An alternate study revealed that elderly mice exhibited a reduced RER compared to their younger counterparts, suggesting that as mice age, there is a corresponding increase in the proportion of fat oxidation [65]. Another study found that, under resting conditions, RER remained comparable between older and younger individuals [66]. It is important to note that RER is just one of many ageing biomarkers that can be used to study biological ageing. RER is influenced by many different factors, such as diet type [67,68], food intake [69], diet restriction [70], extreme ambient temperature, and energy balance [71]. Our study found that PolgA mice show higher food intake (Fig. 5C) compared to their WT counterparts. One of the key nutrients they might be consuming more of is carbohydrates. When carbohydrates are consumed in excess, the body has more readily available energy substrates, leading to increased utilisation of carbohydrates as a fuel source. This increased food intake could contribute to the higher RER observed in PolgA mice. Further, we observed a notch (sharp decline, followed by an increase) in RER in the late active phase in WT mice due to a sharp decrease in food intake (Fig. 5A). In our present study, we did not explore different parameters that influenced the RER index linked to ageing, such as total oxidative metabolism assessment, the association of the RER to physical fitness or endurance, fasting under a controlled temperature, etc. Furthermore, we did not conduct experiments such as constant-dark measurements or feeding-fasting cycles, which could have provided valuable insights into this behaviour within the context of circadian clock regulation. Previous research has demonstrated that food and water intake is influenced more by strain rather than sex differences [72]. We observed that PolgA mice, regardless of sex, displayed a higher propensity for increased food consumption than WT mice at night (Fig. 5A, Fig. 5C). A rise in feeding patterns among mice could show shifts in appetite regulation, metabolic shifts, or age-related physiological transformations. Ageing can influence the control of appetite, energy disbursement, and the use of nutrients. These changes may increase feeding habits, where mice consume more food. Further, increased feeding habits might be due to the PolgA mice showing osteosarcopenia [38], and sarcopenia can induce obesity, diabetes, and other metabolic-associated diseases [73]. We examined the water consumption patterns of these mice, as maintaining adequate hydration can significantly impact bodily functions such as digestion, temperature regulation, and cognitive performance [74,75]. Our results reveal comparable water intake levels in both PolgA and WT mice (Fig. 5B). A decrease in body weight was observed in female PolgA mice compared to their WT counterparts (Fig. 5D). Both humans and mice commonly experience a decline in body weight with age [76]. The reduction in body weight is predominantly attributed to a decrease in lean muscle tissue rather than a fat mass loss. Since muscle is denser than fat, substituting muscle with fat plays a significant role in the overall decrease in body weight. Strikingly, previous studies have demonstrated the presence of osteosarcopenia in aged PolgA mice [38], indicating that physical dysfunction and age-related complications may be contributing factors to the weight loss in these mice during the ageing process. Our findings emphasize the significance of body weight as an essential metabolic parameter to consider when studying the effects of ageing and potential genetic influences. We evaluate the locomotor activities of these mice, as the investigation into age-related alterations in memory, cognitive functions, and motor abilities has been extensively studied in prior research [77–80]. Investigations have shown decreased locomotory behaviour and overall activity in ageing populations [81–83]. These changes are often accompanied by underlying physiological shifts, including sarcopenia or dynapenia in skeletal muscles [38,38,84]. Our analysis unveiled a notable sex difference in total movement, with female WT mice exhibiting higher locomotor activity than their male counterparts (Fig. 6A, Fig. 6B). However, in contrast, female PolgA mice demonstrated diminished walking distance compared to the WT group (Fig. 6C) and walking speed (Fig. 6D). These findings are consistent with previous reports that suggest a decline in locomotor activity in aged animals. Further, we observed that female WT and PolgA exhibit higher energy expenditure than their male counterparts, with female WT demonstrating more energy expenditure than PolgA mice. This outcome implies a tendency for walking speed to diminish in ageing PolgA mice compared to WT mice, aligning with earlier research that demonstrated age-related declines in walking speed [85]. Our analysis also revealed a daily rhythmic pattern in total movement among these mice (Fig. 6A, Fig. S1) and the walking speed (Fig. S2). Grip strength is frequently referred to as a biomarker of ageing, indicative of various factors linked to the decline in strength commonly observed with age. These factors encompass alterations in muscle structure and composition due to ageing, leading to reductions in muscle fiber size and density. Our study showed diminished grip strength in female PolgA mice compared to the wild-type counterparts (Fig. 6E). The reduced grip strength observed in PolgA mice corresponds with their decreased body weight, as indicated in (Fig. 5D). It’s worth noting that weight loss or reduction in body fat can potentially impact grip strength. Given this context, the current findings suggest that female PolgA mice have reached a point where their mobility becomes compromised.

### Conclusions

Our results indicate that PolgA mice exhibit an accelerated ageing phenotype at 41-42 weeks compared to their WT counterparts of the same age. Further, female PolgA mice exhibit a more noticeable ageing compared to their male counterparts. Therefore, PolgA mice can be used as a model to study ageing-related research work efficiently in the future. The home-cage-like platform can be replicated to characterize behaviour, physiology, and intervention testing in different laboratory model organisms. Such study design aligns with Directive 2010/63/EU and the 3R principles (replacement, reduction, refinement). One of the limitations of the present study is that the precise mechanism underlying the ageing effect through metabolic parameters was not explored, and further detailed studies are needed to explore the signalling molecules involved in the metabolic effects. Additionally, the finding relies on metabolic cage measurements of oxygen consumption, carbon dioxide release, locomotor activity, and feeding behaviour in freely moving animals. Therefore, further experiments will be required to explore how ageing mice impact the phase resetting of circadian clocks by changing the light-dark cycle and feeding patterns.

## Acknowledgments

We thank Susanne Friedrich, the Head of Phenotyping at ETH Zurich, for providing valuable support in conducting the phenotypic studies. Funding was provided by European Research Council (ERC Advanced MechAGE ERC-2016-ADG-741883).

## Data availability

The authors confirm that the data supporting the findings of this study are available within the article and its supplementary materials. The raw data can be available from the corresponding author upon request.

## Author Contributions

A.S., E.W., G.K., and R.M. contributed to the design of the study. A.S. conducted the data analysis and visualisation and drafted the manuscript. D.Y. analysed the grip strength. All the authors revised the manuscript collaboratively, and they unanimously approved the final version for submission.

## Abbreviations

VO2: Oxygen consumption
VCO2: carbon dioxide production
RER: Respiratory exchange ratio
LOESS: Locally estimated scatterplot smoothing
ANOVA: Analysis of variance
WT: wild-type littermates
PolgA: PolgA_(D257A/D257A)_
F: Female
M: Male;
EE: Energy Expenditure

## Supporting information

**Table S1:** The table contains the mean and standard deviation of metabolic and locomotory parameters of PolgA and WT mice.

**Table S2:** Analysis of variance results for daily rhythm parameters, body weight, and grip strength, with *p*-values comparing day and night between PolgA and WT mice.

**Fig. S1.** Daily cycle of locomotor activity of PolgA and WT mice.

**Fig. S2:** Daily cycle of walking speed.

