## Supplementary Figure-1 for "Daily rhythms in metabolic and locomotor behaviour of prematurely ageing PolgA mice"

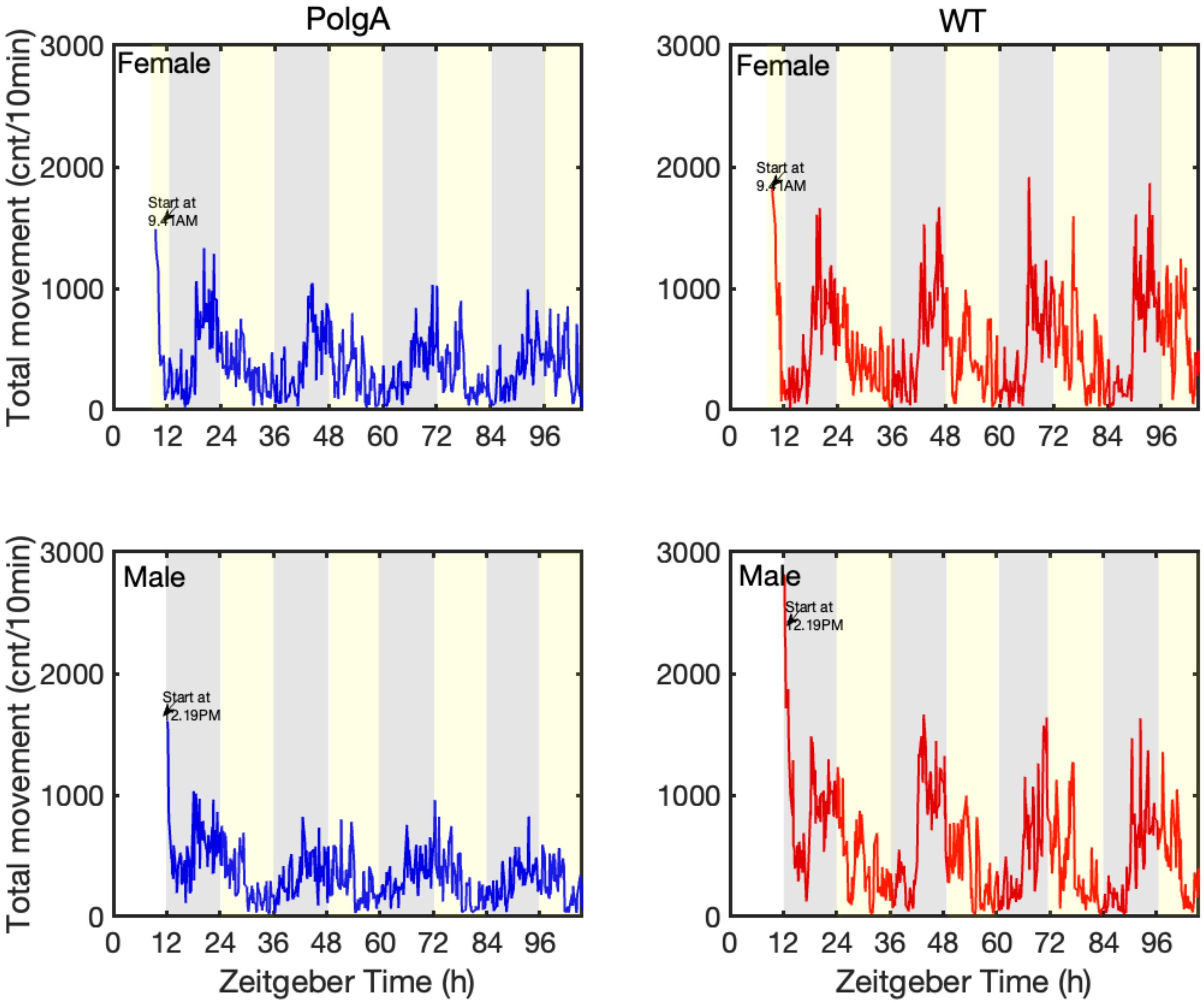


**Supplementary Fig. S1: Daily cycle of locomotor activity of PolgA and WT mice.** The data depict the total walking movements of PolgA (left panel) and WT (right panel) mice, and the measurements were recorded at 10-minute intervals over four consecutive days and nights. The data are presented as average values of locomotor activity (WT male (n=8), WT female (n=7), PolgA male (n=8), and PolgA female (n=8)).
