## Supplementary Figure-2 for "Daily rhythms in metabolic and locomotor behaviour of prematurely ageing PolgA mice"

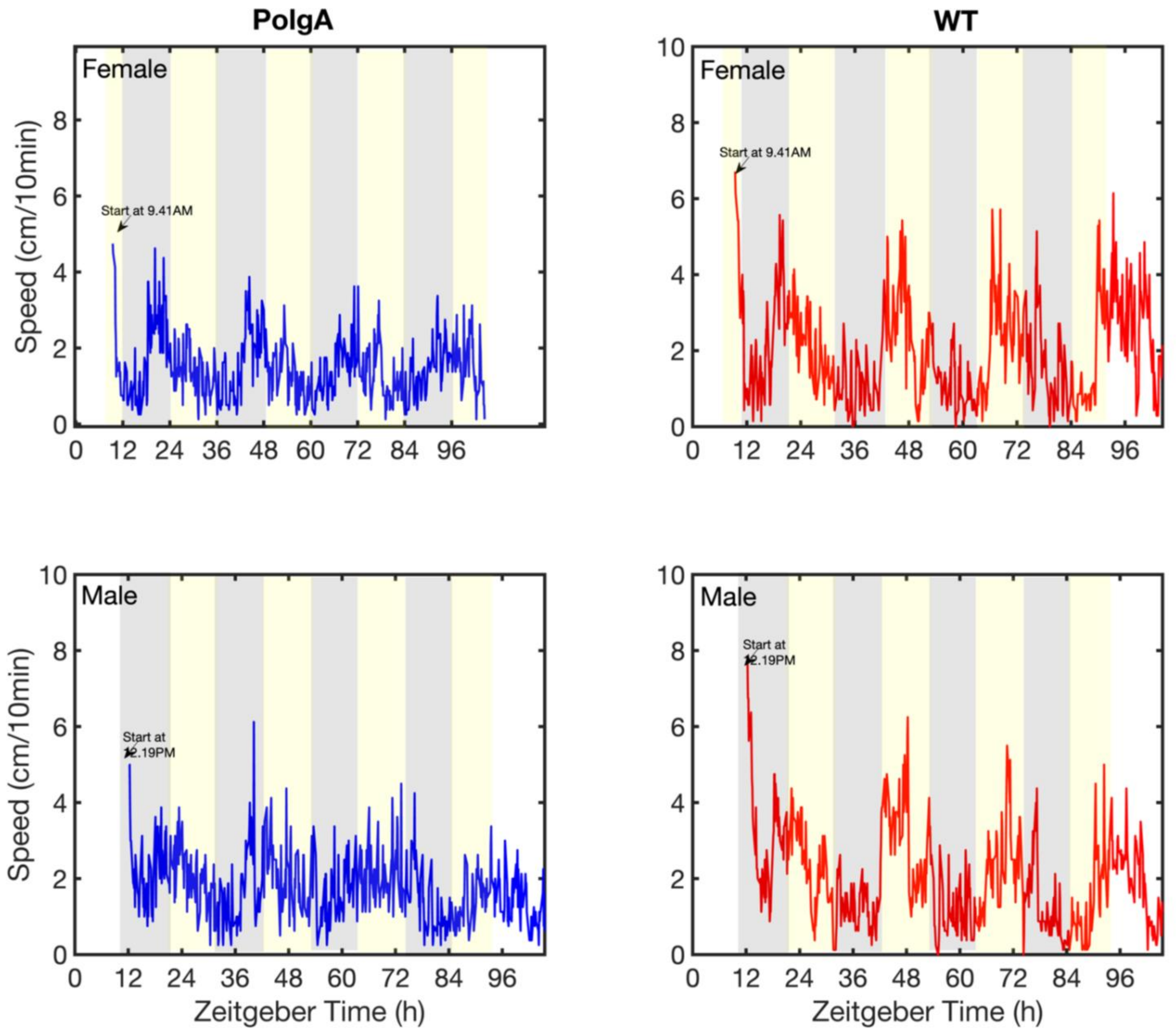


**Supplementary Fig. S2: Daily cycle of walking speed.**

The data illustrate the walking speed of PolgA mice (left panel) and WT mice (right panel) measured in the metabolic cage over four consecutive days and nights. The data are presented as average values of walking speed (WT male (n=8), WT female (n=7), PolgA male (n=8), and PolgA female (n=8)).
